# Proteotoxic stress-induced Nrf1 transcriptional program requires a functional TIP60 complex

**DOI:** 10.1101/443937

**Authors:** Janakiram R. Vangala, Senthil K. Radhakrishnan

## Abstract

In response to inhibition of the cellular proteasome, the transcription factor Nrf1 (also called NFE2L1) induces transcription of proteasome subunit genes resulting in the restoration of proteasome activity and thus enabling the cells to mitigate the proteotoxic stress. To identify novel regulators of Nrf1, we performed an RNA interference screen and discovered that the AAA+ ATPase RUVBL1 is necessary for its transcriptional activity. Given that RUVBL1 is part of different multi-subunit complexes that play key roles in transcription, we dissected this phenomenon further and found that the TIP60 chromatin regulatory complex is essential for Nrf1-dependent transcription of proteasome genes. Consistent with these observations, Nrf1, RUVBL1, and TIP60 proteins were co-recruited to the promoter regions of proteasome genes after proteasome inhibitor treatments. More importantly, depletion of RUVBL1 or TIP60 in various cancer cells sensitized them to cell death induced by proteasome inhibition. Our study provides a framework for manipulating the Nrf1-TIP60 axis to alter proteasome function in various human diseases including cancer.

## INTRODUCTION

Protein homeostasis or proteostasis refers to the sum total of highly regulated and often interconnected cellular processes that include synthesis, folding, and destruction of proteins (1). The ubiquitin-proteasome system (UPS) plays a pivotal role in proteostasis by orchestrating protein degradation. A central component of the UPS is the 26S proteasome, a multi-subunit molecular machine with proteolytic activity that is directly responsible for degrading its protein substrates which are often tagged with polyubiquitin signals. The 26S proteasome is in turn composed of a 20S core, which serves as the main proteolytic chamber, and is capped on one or both ends by the 19S regulatory particle (2). Cellular proteasome activity is regulated at multiple levels including transcription, translation, assembly, and post-translational modifications of subunits (3).

We and others previously demonstrated that the transcription factor Nuclear factor erythroid derived 2-related factor 1 (Nrf1; also called NFE2L1) functions as a master regulator of proteasome subunit genes (4-7). Nrf1 belongs to the cap ‘n’ collar basic-region leucine zipper (CNC-bZIP) family of transcription factors which also includes p45 Nfe2, Nrf2, and Nrf3 (8,9). While Nrf2, the most studied of the CNC-bZIP factors, responds to oxidative stress (10), Nrf1 takes center-stage when cells experience proteotoxic stress (4,6). In response to proteasome inhibition, Nrf1 pathway is activated leading to *de novo* synthesis of proteasome subunit genes, resulting in the “bounce-back” or recovery of proteasome activity (4). Apart from responding to proteotoxic stress, Nrf1 is also responsible for basal transcription of proteasome genes in a tissue-specific manner. Nrf1-deficient mouse neuronal cells display accumulation of ubiquitinated proteins and decreased proteasome activity concomitant with a reduction in proteasome gene expression (11). Similar phenomenon is also evident in hepatocytes from mice that have a liver-specific knockout of Nrf1 (12).

From a molecular perspective, Nrf1 is co-translationally inserted into the endoplasmic reticulum (ER) membrane in such a way that only a small portion of its N-terminus protrudes into the cytosol, whereas a bulk of its polypeptide including the transcriptional activation and DNA-binding domains are embedded in the ER lumen. Thus, Nrf1 depends on the action of the AAA+ ATPase p97/VCP to be retrotranslocated to the cytosolic side either to be constitutively degraded in the absence of proteotoxic stress, or activated and mobilized to the nucleus when cells are subjected to proteasome inhibition (13). Activation of Nrf1 also requires its deglycosylation which is mediated by a p97-associated N-Glycanase enzyme NGLY1 (14) and a proteolytic cleavage step which serves to trim the N-terminus of Nrf1 that harbors the transmembrane domain. The protease involved in this process was recently discovered to be DDI2, a member of the aspartic family of proteases (15,16).

Proteasome inhibitors are currently being used in the clinic against multiple myeloma and mantle cell lymphoma (17). One way to make this therapy more effective would be to devise strategies to block the Nrf1 pathway, thus impairing the ability of the cancer cells to recover their proteasome activity in response to inhibition of the proteasome. In support of this notion, a recent study demonstrated that attenuation of Nrf1 activation via p97 inhibition increased the efficacy of proteasome inhibitor drugs in multiple myeloma models (18). On the flip side, in certain proteinopathies, especially neurodegenerative diseases where the neurons display ubiquitinated protein aggregates indicative of proteasome dysfunction, there could be interest in enhancing the Nrf1 pathway to directly increase proteasomal capacity (3). Thus, further understanding of the Nrf1-proteasome axis could help pave the way for devising novel therapeutics targeted at different human diseases. In this study, we identify a requirement for the TIP60 chromatin-modifying complex in enabling Nrf1-mediated proteasomal gene expression in response to inhibition of the proteasome. This expands our view of the inner workings of the Nrf1-mediated transcription and could offer strategies to manipulate this pathway in human diseases.

## RESULTS

### RUVBL1 is necessary for Nrf1-mediated transcriptional response during proteotoxic stress

To identify factors that regulate Nrf1 activity under conditions of proteasome inhibition, we constructed a cell-based screening system in wild-type (WT) NIH-3T3 mouse fibroblasts. This cell line was engineered to stably express firefly luciferase under the control of 8× anti-oxidant response element (ARE; the sequence to which Nrf1 is known to bind (8)) repeats coupled to a minimal promoter. In addition, as a control, these cells also expressed renilla luciferase driven by human phosphoglycerate kinase (hPGK) promoter (Fig 1A). The resultant screening system is referred to as WT 8×ARE-Luc, whereas a similar system in NIH-3T3 Nrf1-deficient background (19) is referred to as Nrf1^-/-^ 8xARE-Luc. We then confirmed the status of Nrf1 in these two cell lines. Treatment with proteasome inhibitor carfilzomib (CFZ) resulted in the accumulation of Nrf1 p120 (precursor) and p110 (processed form; transcriptionally active) in the WT 8xARE-Luc but not in Nrf1^-/-^ 8xARE-Luc cell line as expected (Fig 1B). More importantly, whereas the WT 8xARE-Luc cells showed a dose-dependent increase in normalized luciferase activity (ratio of firefly to renilla luciferase values) in response to CFZ, the Nrf1^-/-^ 8xARE-Luc cell line showed no such increase (Fig 1C), implying that this effect is Nrf1-dependent. In light of our previous study that demonstrated a strict reliance of Nrf1 function on p97 ATPase activity (13), we used NMS-873, an inhibitor of p97 (20) to further test our screening system. NMS-873 was able to effectively attenuate CFZ-induced increased luciferase activity in WT 8xARE-Luc cell line (Fig 1D), thus additionally validating our screening system.

**Figure 1.**
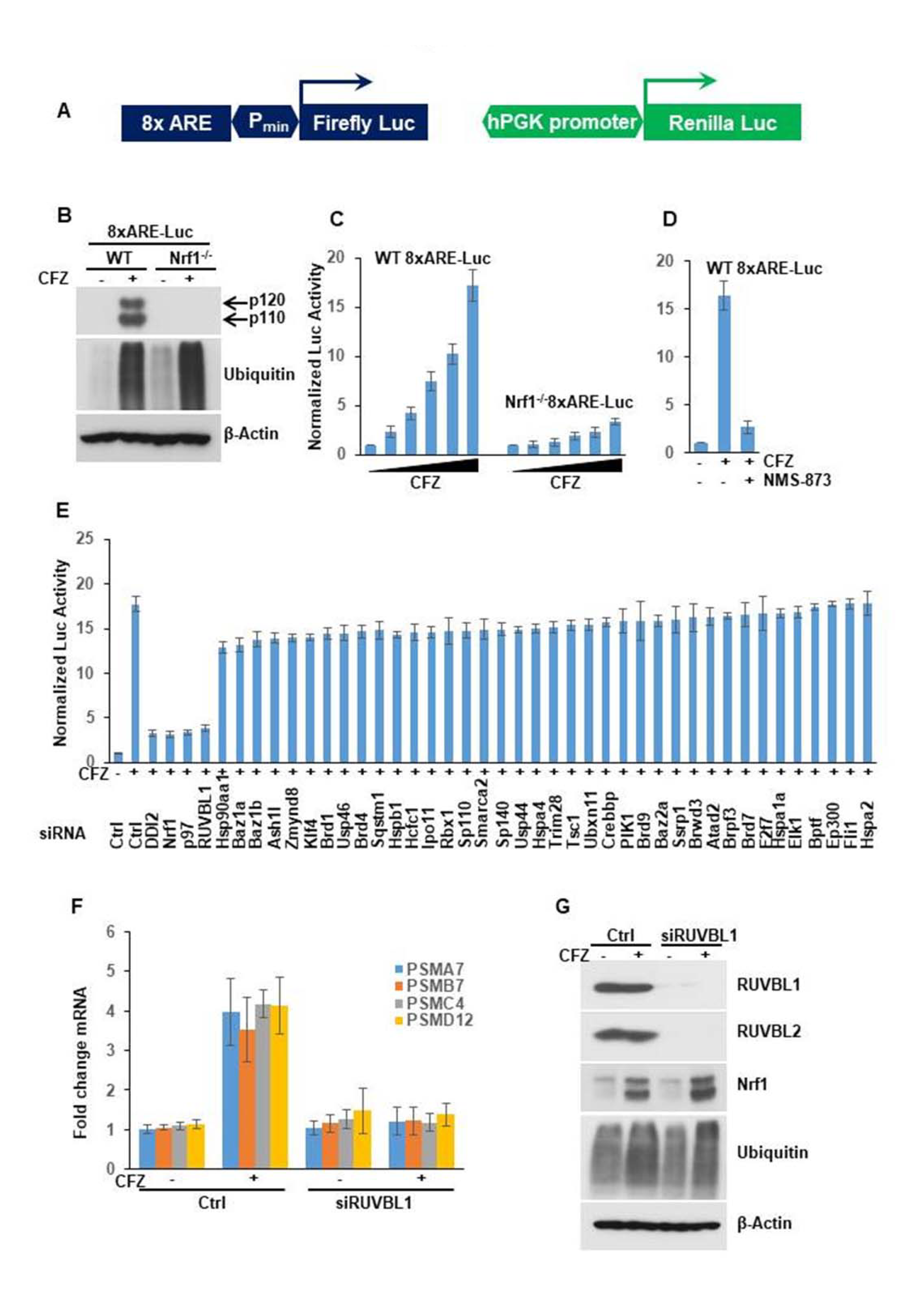
Identification of RUVBL1 as a factor that is required for Nrf1 transcriptional activity. **(A)** Schematic representation of the reporter construct used in the cell-based screening system is shown. This lentiviral reporter construct expressed firefly luciferase under the control of 8x antioxidant response elements (8xARE) upstream of a minimal promoter (P_min_), along with renilla luciferase driven by hPGK promoter. **(B)** NIH-3T3 cells with stable incorporation of the above described reporter system and that were either wild-type (WT 8xARE-Luc) or Nrf1-deficient (Nrf1^-/-^ 8xARE-Luc) were treated with DMSO or 200 nM carfilzomib (CFZ) overnight and analyzed by immunoblotting using antibodies specific for Nrf1, ubiquitin, and β-Actin. **(C)** WT 8xARE-Luc and Nrf1^-/-^ 8xARE-Luc cells were treated for 16 hours with increasing concentrations of CFZ (0, 20, 50, 100, 150, 200 nM) and then subjected to Dual Luciferase assays to measure the firefly and renilla luciferase activity values. Normalized luciferase activity is shown. Error bars denote SD (n=3). **(D)** WT 8xARE-Luc cells were treated with 200 nM CFZ alone or in combination with 10 μM NMS-873 and compared with DMSO treated control for 16 hours. The cell lysates were then used for luciferase assays. Normalized luciferase activity is shown. Error bars denote SD (n=3). **(E)** WT 8xARE-Luc cells were transfected with a focused library of siRNAs targeting several epigenetic factors and other candidate genes. Forty-eight hours after transfection, the cells were further treated with 200 nM CFZ overnight and assayed for luciferase activity. Error bars denote SD (n=3). **(F)** Wild-type NIH-3T3 cells were either control transfected or with siRNAs targeting RUVBL1 and further treated with 200 nM CFZ as indicated for 8 hours. RNA extracted from these cells was then subjected to quantitative RT-PCR with primers specific for representative proteasome subunit genes as shown. The transcript levels of 18S rRNA were used for normalization. Error bars denote SD (n=3). **(G)** NIH-3T3 cells treated as described in (F) were used for immunoblotting with antibodies against Nrf1, RUVBL1, RUVBL2, ubiquitin, and β-Actin as indicated.

Next, we used our WT 8xARE-Luc cell system in a RNA interference (RNAi) screen where we used a focused library of siRNAs that target various epigenetic regulators and possible Nrf1 pathway influencers gleaned from public databases and literature (8,9,21,22). As positive controls, we used Nrf1-specific siRNAs and also siRNAs targeting known Nrf1 regulators such as p97 and the protease DDI2. In the screen, we found that depletion of any of the controls – Nrf1, p97, or DDI2 – strongly attenuated CFZ-induced increase in luciferase activity (Fig 1E). Out of the other test genes in the library, we observed that knockdown of RUVBL1 elicited an effect that was quite similar to the ones produced by the three positive control siRNAs (Fig 1E). The results from the entire screen are listed in Supplemental Table S1.

RUVBL1 (Pontin/TIP49) and the related protein RUVBL2 (Reptin/TIP48) are AAA+ ATPases that most often occur together and participate in diverse cellular functions including transcriptional regulation (23). Given that we identified RUVBL1 as a ‘hit’ in our screen, it most likely is involved in Nrf1-mediated transcriptional activation of its target genes. To further confirm the involvement of RUVBL1 in the Nrf1 pathway, we examined the changes in proteasome gene transcription in response to CFZ treatment in control and siRUVBL1-treated NIH3T3 cells. Whereas the control cells displayed a functional Nrf1 pathway by upregulating representative proteasome subunit genes as expected, the cells with RUVBL1 depletion were profoundly defective in this response (Fig 1F). Under these circumstances, we did not see a significant difference in Nrf1 protein levels or its ability to be processed into p110 form after RUVBL1 knockdown (Fig 1G). Taken together, these results reinforce a model where RUVBL1 could regulate the transcriptional activity of Nrf1 as has been seen for certain other transcription factors such as c-Myc, β-catenin, E2F1, NF-κB, and HIF1α (23). Interestingly, in the control immunoblots to verify knockdown efficiency in this experiment, we noticed that apart from RUVBL1, the protein levels of RUVBL2 were also attenuated in lysates derived from siRUVBL1-treated samples (Fig 1G). This is consistent with previous observations where depletion of either factor results in a concomitant decrease in the other, in line with the notion that RUVBL1 and RUVBL2 together need to be incorporated into a hexameric complex to attain stability (24-26).

To rule out any cell type-specific bias, we tested other cell lines of various origin – HCT116 colon cancer, MDA-MB-231 breast cancer, and MIA-PaCa2 pancreatic cancer cells. Regardless of the cell type, we observed that whereas proteasome inhibition upregulated the transcript levels of proteasome genes tested, depletion of RUVBL1 (Fig S1) severely compromised this effect (Fig 2A). Similar to our previous observation with NIH-3T3, in all of these cancer cell lines, depletion of RUVBL1 resulted in decrease in the protein levels of both RUVBL1 and RUVBL2 (Fig 2B).

**Figure 2.**
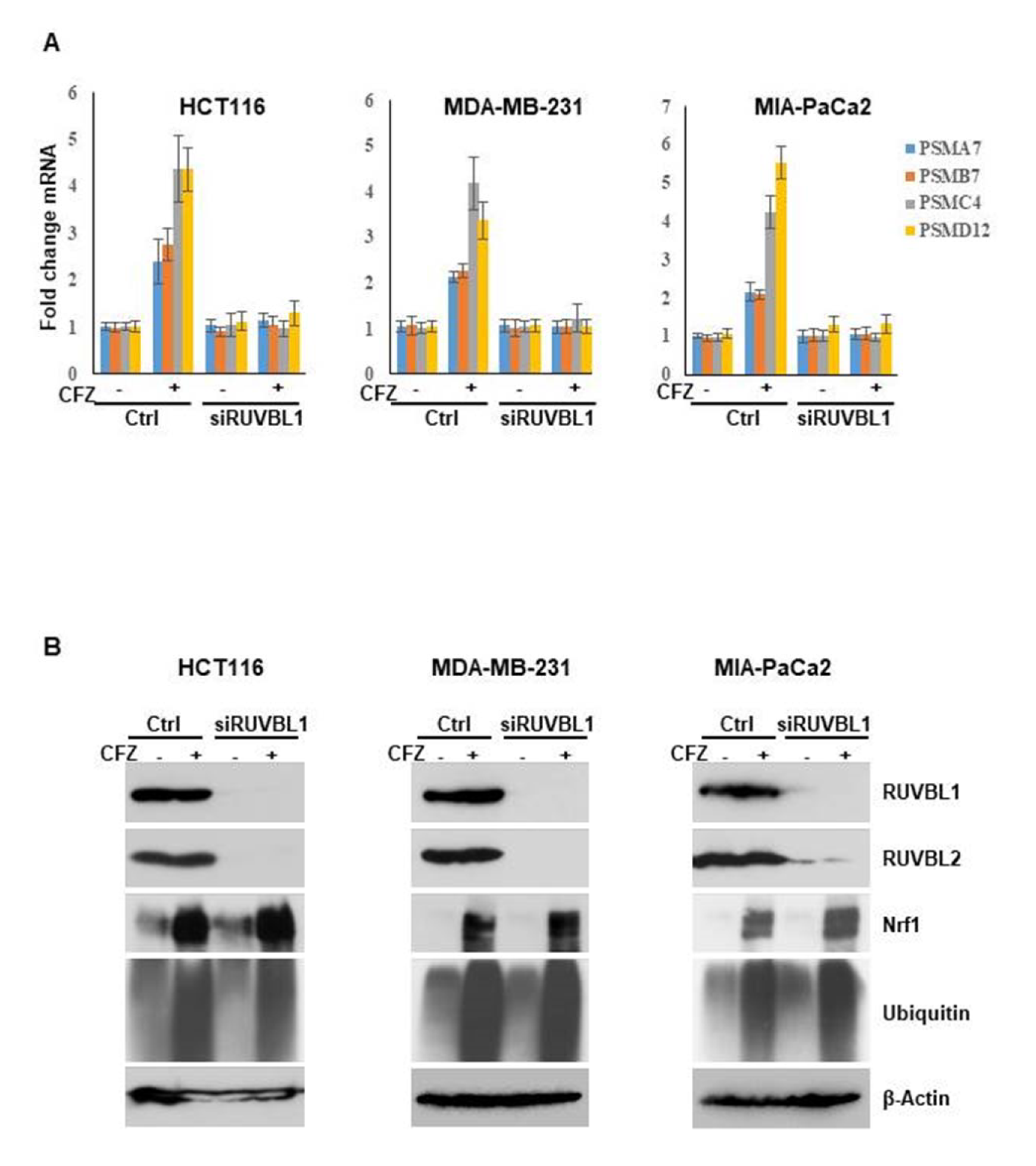
Depletion of RUVBL1 impairs the transcriptional function of Nrf1 in different cancer cell lines. **(A)** The cell lines HCT116, MDA-MB-231, and MIA-PaCa2 were either control transfected or with siRNAs targeting RUVBL1. Forty-eight hours after transfection, the cells were treated with 200 nM carfilzomib (CFZ) for 8 hours and then analyzed by quantitative RT-PCR to measure representative proteasome subunit gene mRNA levels. The mRNA levels of 18S rRNA were used for normalization. Error bars denote SD (n=3). **(B)** The cell lines above were treated similarly as described in (A) and subjected to immunoblotting with antibodies specific for RUVBL1, RUVBL2, Nrf1, ubiquitin, and β-Actin.

### TIP60 complex is required for Nrf1-dependent transcription

Although in some instances, RUVBL1 and RUVBL2 act as a heterohexamer to regulate gene transcription (27), in most cases these proteins are part of multi-subunit complexes with roles in transcription (23). The most prominent RUVBL1/2-containing complexes are INO80, SRCAP, TIP60, and R2TP (Fig 3A). To test, if one or more of these complexes could be involved in mediating Nrf1-dependent transcription under proteotoxic stress, we proceeded to deplete a key component of each complex (INO80 subunit in the INO80 complex, SRCAP subunit in the SRCAP complex, TIP60 subunit in the TIP60 complex, and PIH1 subunit in the R2TP complex) in the WT 8xARE-Luc cell line. We observed that depletion of TIP60, but not any of the other subunits tested (Fig S2), phenocopied the effect of RUVBL1 knockdown in blocking the proteasome inhibitor-mediated increase in luciferase activity (Fig 3B). Likewise, when we measured transcriptional upregulation of proteasome genes in response to proteasome inhibition, depletion of TIP60 attenuated this response similar to what was observed with RUVBL1 depletion (Fig 3C). Taken together, our results attribute a critical role for the TIP60 complex in Nrf1-dependent transcriptional response to proteotoxic stress. Interestingly, in our control immunoblots, we observed that depletion of RUVBL1 also decreased TIP60 and to a certain extent INO80 and PIH1 protein levels (Fig 3D). This is consistent with an earlier study that indicated RUVBL1/2 complex is essential for the assembly of a functional TIP60 complex (28).

**Figure 3.**
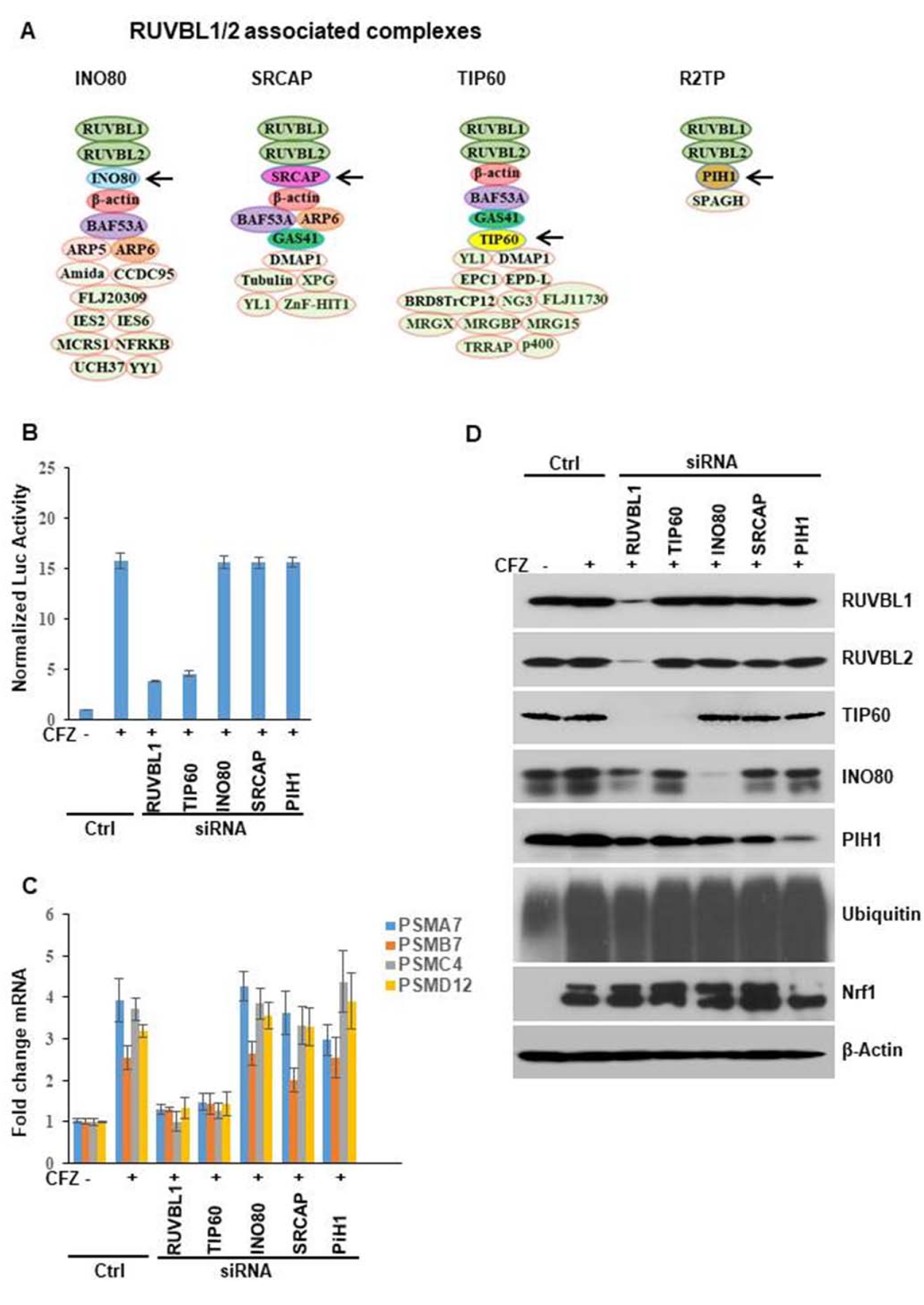
TIP60 complex is required for Nrf1 transcriptional activity. **(A)** Some of the multi-subunit complexes that contain RUVBL1 are depicted. The subunits shown in red are targets of siRNAs in the panels below. **(B)** WT 8xARE-Luc cells were transfected with siRNAs targeting RUVBL1, TIP60, INO80, SRCAP, and PIH1 as indicated. Forty-eight hours after transfection, the cells were further treated with 200 nM carfilzomib (CFZ) overnight and luciferase assays were performed. Normalized luciferase activity is shown. Error bars denote SD (n=3). **(C)** Wild-type NIH-3T3 cells were transfected and further treated as described in (B) and subjected to quantitative RT-PCR to measure mRNA levels of select proteasome subunit genes. The mRNA levels of 18S rRNA were used for normalization. Error bars denote SD (n=3). **(D)** Wild-type NIH-3T3 cells were treated as described above in (C) and the lysates were used for immunoblotting to measure the levels of different proteins as indicated. β-Actin was used as a loading control.

### Interaction between Nrf1 and the TIP60 complex

Based on the known biology of TIP60 as a transcriptional co-activator, it is possible that this complex interacts with Nrf1 on the chromatin to aid in its transcription function. To verify this hypothesis, we compared the ARE-containing promoter regions of Nrf1 target genes PSMA7, PSMB7, and PSMD12 in WT and Nrf1^-/-^ NIH3T3 cells using chromatin immunoprecipitation (ChIP). Under the conditions of proteasome inhibition with CFZ, we could observe recruitment of Nrf1 in the promoter regions of proteasome genes in the WT, but not Nrf1^-/-^ cells as expected (Fig 4A). Importantly, we observed a similar trend for RUVBL1 and TIP60 subunits implying Nrf1-dependent recruitment of these factors to the proteasome gene promoters. Consistent with this notion, we were also able to detection interaction between Nrf1 and RUVBL1/2 using co-immunoprecipitation assays (Fig 4B).

**Figure 4.**
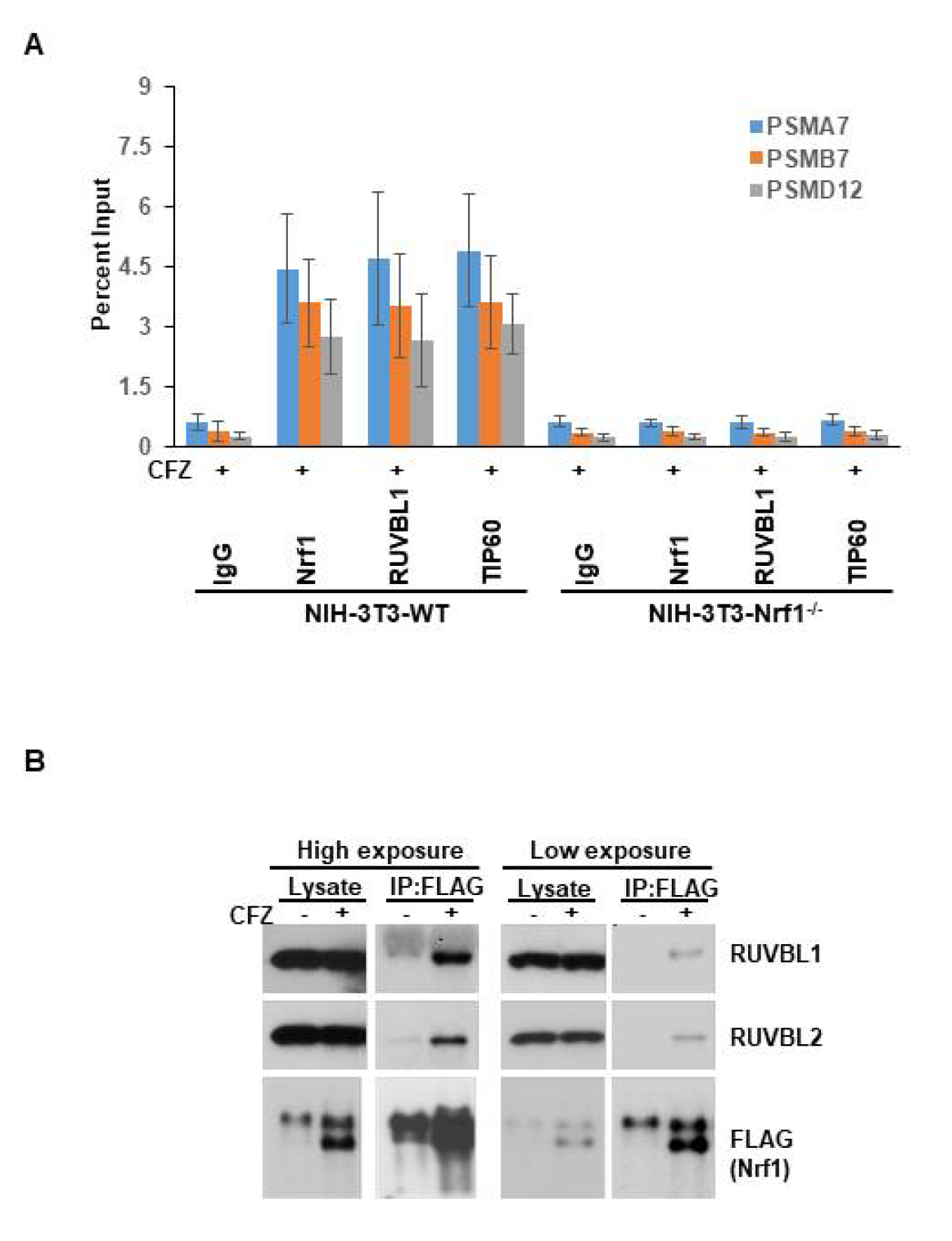
Nrf1 interacts with the TIP60 complex. **(A)** Wild-type (WT) and Nrf1^-/-^ NIH-3T3 cell lines were treated with 200 nM carfilzomib (CFZ) for 8 hours. The cells were then subjected to chromatin immunoprecipitation (ChIP) with one of IgG, Nrf1, RUVBL1, or TIP60 antibodies. These samples were then analyzed by quantitative PCR with primers specific for antioxidant response element (ARE)-containing promoter regions of proteasome genes PSMA7, PSMB7, and PSMD12. **(B)** HEK293 cells stably expressing tagged Nrf1 (Nrf1^3xFLAG^) were treated or not with 200 nM CFZ for 8 hours. The cell lysates were then subjected to immunoprecipitation with anti-FLAG beads and analyzed by immunoblotting with antibodies specific for FLAG, RUVBL1, and RUVBL2.

### Functional consequences of depletion of TIP60 complex during proteotoxic stress

Our previous work conclusively demonstrated a cellular requirement for Nrf1 in recovering from inhibition of proteasome (4). To verify if this effect is recapitulated in cells deficient in RUVBL1 or TIP60, we performed proteasome recovery assays (Fig 5A). Here, we exploited the differences in the nature of binding of proteasome inhibitors to the active site of the proteasome. Whereas MG132 is a completely reversible proteasome inhibitor, CFZ binds to the proteasome active site in a covalent/irreversible manner. When cells are treated with MG132 briefly and then washed off, the recovery of the proteasome activity is achieved by simple dissociation of the drug from the proteasome active site, as well as via the action of the Nrf1 pathway during the time when the proteasome was inhibited. In the case of CFZ, given its covalent binding to the proteasome, the only way for the cells to recover after drug wash-out is by invoking the Nrf1 pathway to synthesize new proteasomes. Thus, impairing Nrf1 function in this context will severely undermine the ability of the cells to recover proteasome activity (4). To test these predictions in our study, we compared wild-type NIH-3T3 cells with siRUVBL1 or siTIP60 transfected cells for their relative ability to recover proteasome activity after pulse-treatment of MG132 or CFZ with appropriate concentrations to achieve about 90% inhibition of proteasome activity in one hour. Following drug wash-out, we found that, cells depleted of RUVBL1 or TIP60 were able to recover their proteasome activity after MG132 pulse-treatment, but were severely impaired in their ability to do so when CFZ was employed (Fig 5B). These results suggest that depletion of RUVBL1 or TIP60 phenocopy Nrf1-loss in the context of proteasome recovery.

**Figure 5.**
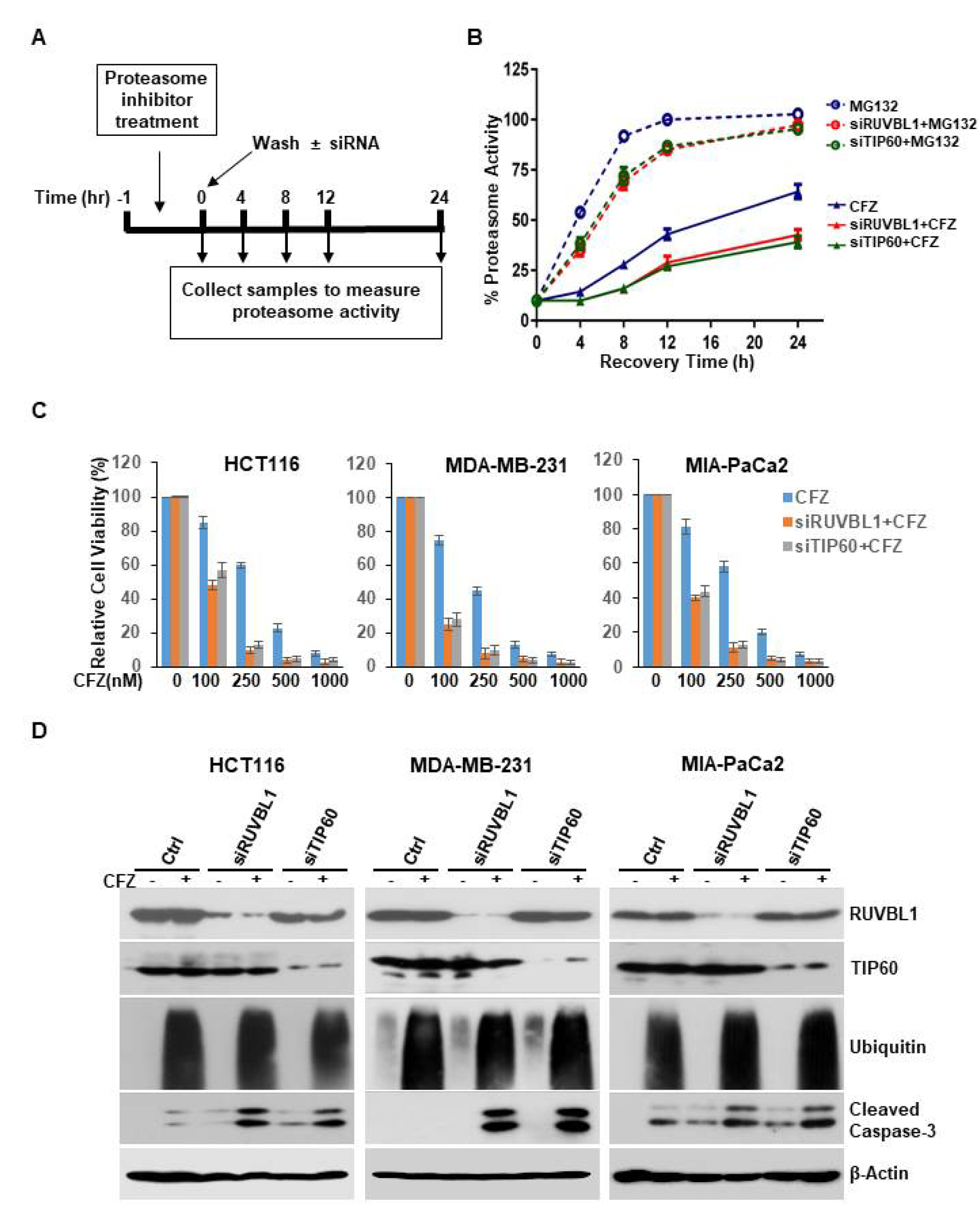
Depletion of RUVBL1 or TIP60 impairs proteasome recovery and potentiates carfilzomib-mediated apoptosis. **(A)** Schematic representation of the proteasome recovery assay is shown. **(B)** NIH-3T3 cells were control transfected or with siRNAs specific for RUVBL1 or TIP60. The cells were then treated for an hour with either 500 nM MG132 or 50 nM carfilzomib (CFZ). The drugs were then washed out, and proteasome activity in the lysates was measured at indicated time points. The results were normalized to DMSO-treated control. Error bars denote SD (n=3). **(C)** The cell lines HCT116, MDA-MB-231, and MIA-PaCa2 were either control transfected or with siRNAs targeting RUVBL1, and TIP60 as indicated. Forty-eight hours after transfection, the cells were treated with increasing concentrations of CFZ for further 24 hours. The cells were then subjected to viability assays. **(D)** The different cancer cell lines were transfected and as described in (A) and then further treated with 250 nM CFZ for 24 hours. The cell lysates were then used for immunoblotting to measure protein levels of cleaved caspase-3 along with levels of RUVBL1, TIP60, ubiquitin, and β-Actin as controls.

Irresolvable proteotoxic stress could lead to cell death, particularly in cancer cells which are thought to be over-reliant on proteasome function (29,30). Our previous studies have demonstrated potentiation of apoptosis in Nrf1-impaired cancer cells that were treated with proteasome inhibitors (4,14). To evaluate the contribution of the TIP60 complex in this context in cancer cells, we first exposed control and siRUVBL1 or siTIP60 treated cells (HCT116, MDA-MB-231, and MIA-PaCa2) to different concentrations of CFZ. While CFZ by itself caused a dose-dependent decrease in cell viability, this effect was exacerbated in siRUVBL1 and siTIP60 treated cells (Fig 5C). Consistent with these results, immunoblotting experiments revealed that when compared to control, the levels of cleaved caspase-3 (a marker for apoptosis) were markedly elevated in RUVBL1 or TIP60 depleted cells that were further treated with CFZ (Fig 5D). Taken together, our results indicate that modulating the activity of TIP60 complex could enable manipulation of the functional output from the Nrf1 pathway in cells experiencing proteotoxic stress.

## DISCUSSION

Nrf1 has emerged as a critical transcription factor in the cellular arsenal to combat proteotoxic stress or proteasome insufficiency. In response to inhibition of the proteasome, Nrf1 is mobilized to enable increased transcription of proteasome subunit genes culminating in the formation of new proteasomes (4,6,13,19). Several previous studies, including ours, have dissected the regulation of Nrf1 function that is achieved via multiple mechanisms in the cell – regulation of its abundance by ubiquitin ligases HRD1, FBXW7, and β-TRCP; regulation of its extraction from the endoplasmic reticulum (ER) by p97; regulation of its proteolytic processing by DDI2; regulation of its deglycosylation by NGLY1; regulation of its phospohorylation and thereby its transcriptional activity by casein kinase 2 – to name a few (9,14,15). Our current study adds the TIP60 chromatin-modifying complex to the growing list of factors that impact Nrf1 activity (Fig 6). To our knowledge, this is the first example of an epigenetic regulator that appears to modulate Nrf1 function.

**Figure 6.**
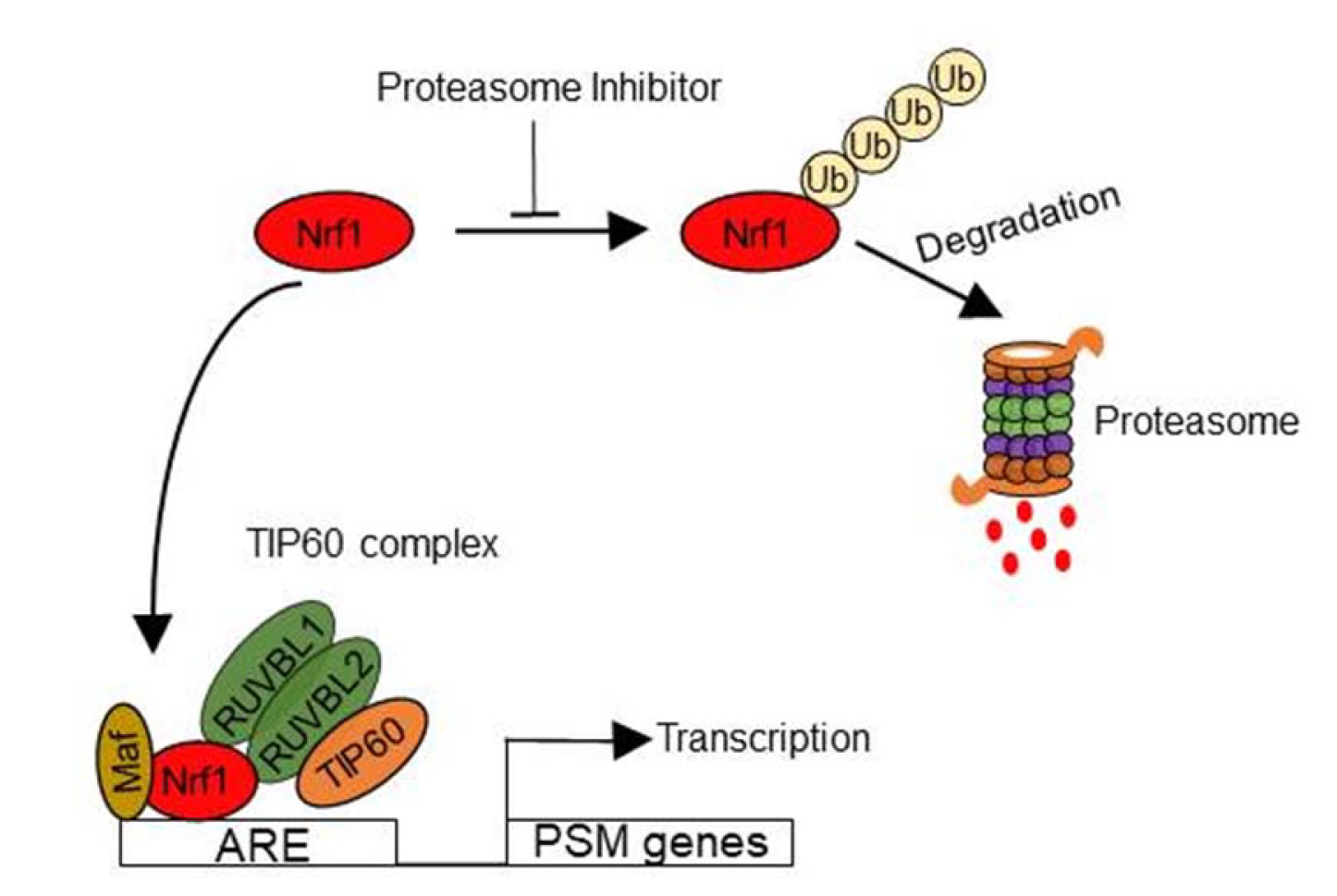
A model to explain the involvement of the TIP60 complex in the Nrf1 pathway. Nrf1 is typically an unstable protein that is constantly ubiquitinated and subjected to degradation by the proteasome. However, if the proteasome is inhibited, Nrf1 is no longer degraded, but is mobilized to the nucleus where together with a small Maf protein binds to the anti-oxidant response elements (ARE) found in the promoter region of its targets that include proteasome subunit (PSM) genes. Based on the known function of the TIP60 complex as a chromatin regulator, we propose that this complex could act as a co-activator of Nrf1.

Although we have demonstrated that RUVBL1 and TIP60, both of which are subunits of the TIP60 complex, are necessary for Nrf1-mediated transcription after proteasome inhibition, the exact nature of their requirement remains to be uncovered. Based on the well-established role for RUVBL1/2 in chromatin remodeling, it could be that these factors serve to open up the chromatin at Nrf1 target gene promoters, as has been seen previously in the case of E2F1 transcription factor (31). Also, TIP60, which harbors acetyl transferase activity, is known to acetylate histones H4 and H2A, thereby relaxing the chromatin in promoter regions and providing access to the transcriptional machinery (32). This is exemplified in the case of hypoxia response where HIF1α recruits TIP60 which enables acetylation of histones and subsequent recruitment and activation of RNA polymerase II at HIF1α target gene promoters (33). In our case, it could be that TIP60 provides such a function in Nrf1-dependent transcription. Given that TIP60 can also directly acetylate some transcription factors such as c-myc and androgen receptor and modulate their function (34,35), it could also be that Nrf1 is subject to such modification. It is important to note that all of these mechanisms may also occur together and are not necessarily mutually exclusive.

Mechanism aside, our findings could have important therapeutic implications, especially in cancer. RUVBL1 and RUVBL2 have been shown to be overexpressed in a number of cancer types including colorectal, liver, breast, lung, gastric, esophageal, pancreatic, kidney and leukemia (23). More importantly, depletion of RUVBL1/2 seemed to delay or even shrink the tumors in preclinical models of some of the cancer types mentioned above. Accordingly, there is intense interest in developing small molecule inhibitors of the ATPase activity of RUVBL1/2 (36,37). These early stage inhibitors already show promise in cell culture and xenograft studies in mice. Likewise, TIP60 has also been regarded as a viable target in cancer and there have been efforts directed to identify inhibitors of its acetyl transferase activity (38). Given that in our study, depletion of RUVBL1 or TIP60 enhances cell killing by carfilzomib, this provides a framework for testing RUVBL1 or TIP60 inhibitors in combination with proteasome inhibitors in various cancer types. This combination would also make sense from the point of view of proteasome inhibitors, which apart from their current use in the clinic against multiple myeloma and mantle cell lymphoma, are now being evaluated in combination with other chemotherapeutic agents in several types of solid tumors (39). Our findings that demonstrates a strong reliance of the Nrf1-mediated proteasome recovery pathway on functional TIP60 complex prompts further consideration of combination trials with proteasome and TIP60 complex inhibitors.

## EXPERIMENTAL PROCEDURES

### Screening system

The lentiviral construct PLKO-luc-mcherry-puro-renilla (Addgene plasmid# 29783) was a gift from Dr. Carl Novina and has been previously described (40). This construct was digested with AgeI and EcoRI to generate PLKO-luc-mcherry-puro. The Renilla luciferase-Neomycin cassette hRluc-neo was PCR-amplified from pF9A CMV hRluc-neo Flexi (C9361; Promega) using primers containing XmaI and EcoRI. The XmaI/EcoRI-digested PCR product was ligated into PLKO-luc-mcherry-puro to generate PLKO-luc-mcherry-puro-renilla-neo. This construct was further digested with SacII and BamHI to release the hPGK promoter driving the luc-mcherry-puro cassette and ligated with 8x antioxidant response element (ARE)-minimal promoter containing oligos harboring overhangs of SacII and BamHI to construct PLKO-8xARE-Pmin-luc-mcherry-puro-hPGK-renilla-neo (refered to as 8xARE-Luc). Whereas this construct expressed firefly and renilla luciferase products as measured by luciferase assays, mcherry expression was undetectable by fluorescent microscopy. Using 8xARE-Luc together with appropriate helper plasmids, lentiviral particles were produced as described previously and used to infect wild-type and Nrf1^-/-^ NIH-3T3 cells. These cells were selected in geneticin and the resultant cell lines are referred to as WT 8xARE-Luc and Nrf1^-/-^ 8xARE-Luc in the other sections of this paper.

### Cell culture and siRNA transfections

All cell lines were grown in Dulbecco’s modified Eagle’s medium (DMEM) supplemented with 10% fetal bovine serum (Atlanta Biologicals), penicillin and streptomycin (Invitrogen) at 37°C in a humidified incubator with 5% CO_2_. All siRNAs (sequences shown in Tables S2 and S3) were purchased from Dharmacon and were transfected into the cell lines using DharmaFECT reagent (Dharmacon) according to manufacturer’s recommendations. Forty-eight hours after transfection, cells were used for downstream experiments.

### Quantitative reverse transcription PCR

TRIzol reagent (Life Technologies) was added directly to the cell culture plates and total RNA was isolated according to manufacturer’s instructions. cDNA was then prepared from 1000 ng of total RNA using iScript cDNA synthesis kit (Bio-Rad). Quantitative PCR was carried out with iTaq universal SYBR green supermix (Bio-Rad) using C1000 Touch Thermal cycler (Bio-Rad). Data was analyzed using CFX manager 3.1 (Bio-Rad). The levels of 18S rRNA were used for normalization. Primers used for the assays are listed in Table S4.

### Immunoblot analysis

Cells were washed twice with ice-cold PBS and pellets were collected under cold conditions. To obtain lysates, cell pellets were resuspended in standard RIPA lysis buffer and incubated on ice for 30 min followed by high speed centrifugation at 13,200 rpm for 30 min. Total protein was quantified by Bradford reagent. Typically, 30 μg of protein was used for SDS-PAGE, followed by electro-transferring on to polyvinylidene difluoride membranes. The membranes were then blocked for 1 hour with 5% non-fat dry milk powder in Tris-buffered saline with Tween and then incubated with the appropriate primary antibodies. The antibodies used were specific for Nrf1, RUVBL1, RUVBL2, Ubiquitin, cleaved caspase-3 (all from Cell Signaling), INO80 (a gift from Dr. Landry), PIH1, (Proteintech) and TIP60 (Abcam) and β-Actin (Sigma-Aldrich). The secondary antibodies used were rabbit IgG HRP, and mouse IgG HRP (both from Bio-Rad).

### Chromatin Immunoprecipitation

Ten-million cells were fixed with freshly prepared Formaldehyde solution (1% final volume) and incubated at room temperature for 10 min, followed by the addition of Glycine to quench the formaldehyde. Plates were then washed twice with ice cold PBS and collected in PBS supplemented with protease inhibitor cocktail. Pellets were collected after centrifugation at 800g at 4 °C. EZ-Magna ChIP A/G (Millipore) kit was used for further steps according to manufacturer’s protocol. Briefly, cells were lysed in cell lysis buffer and the nuclei were pelleted. These pellets were suspended in nuclear lysis buffer and used for shearing chromatin with Covaris M220. Shearing was done at 10% df for 16 min. Sheared chromatin was centrifuged at 10,000g for 10 min and supernatant was collected. 20 μL of protein A/G magnetic beads were used for pre-clearing the sheared chromatin for 1 hr at 4 °C on rotor. 50 μL of pre-cleared chromatin was used for each immunoprecipitation with 5 μg of specific antibody for overnight at 4 °C. Beads were then washed with low, high salt, LiCl, followed by TE buffer. Elution was done with elution buffer at 65 °C for 4 hours followed by column purification (QIAGEN) to obtain purified chromatin. qPCR was used to analyze the chromatin. Primers used for analysis are shown in Table S4.

### Proteasome activity recovery assays

Cells grown in 96-well plates were treated for an hour with 500 nM MG132 or 50 nM carfilzomib, the doses that were previously determined to inhibit the proteasome activity by 90%. The cells were then washed with PBS thrice and allowed to recover in fresh medium. The cells were frozen in TE buffer at different time points. At the time of the assays, the cells were thawed and used for measuring proteasome activity as described previously (4).

### Co-immunoprecipitation assays

HEK293 cells stably expressing C-terminally tagged Nrf1 (Nrf1^3xFLAG^) after appropriate treatments were lysed in IP buffer (50 mM Tris pH 7.4, 0.5 M NaCl, 1 mM EDTA, 1% Triton X-100) supplemented with protease and phosphatase inhibitor cocktail (Pierce). The lysates were then used for immunoprecipitation with anti-FLAG beads (Sigma-Aldrich) as per manufacturer’s recommendations. The samples were eluted in Laemmli buffer and analyzed by SDS-PAGE followed by immunoblotting.

### Cell viability assays

Cells were typically grown in 96-well plates and used for quantifying viability with Cell-Titer Glo kit (Promega) that measures the level of ATP which in turn is proportional to the number of viable cells. Qualitative measure of cell viability was assessed using immunoblots for cleaved caspase-3.

## ACKNOWLEDGEMENTS

We thank A. Feygin for technical assistance. S.K.R. was supported by a K99/R00 award from the National Cancer Institute (R00CA154884) and from a grant from the Grace Science Foundation.

## Supporting Information

**Figure S1.**
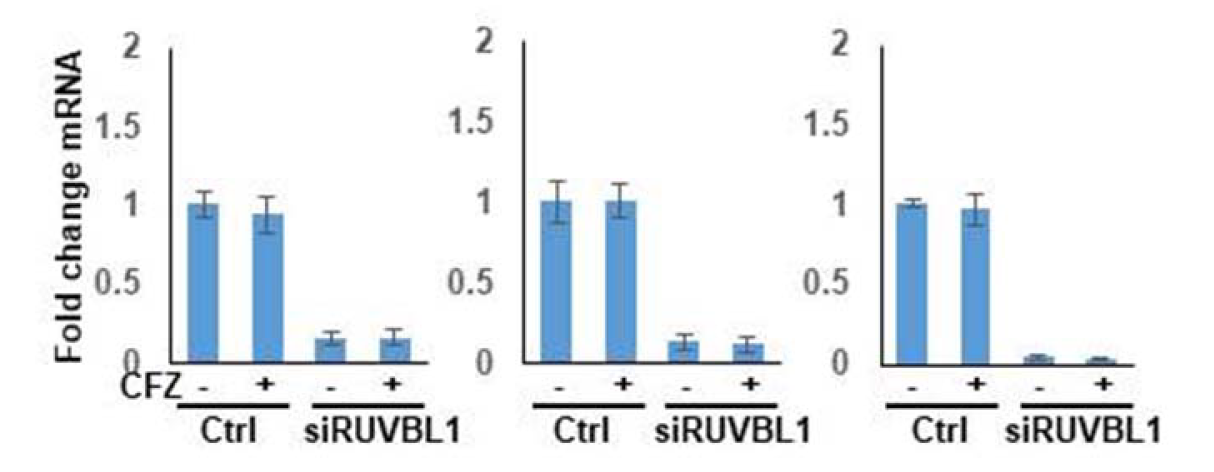
Knock-down efficiency of siRNAs targeting RUVBL1. The cell lines HCT116, MDA-MB-231, and MIA-PaCa2 were either control transfected or with siRUVBL1. Forty-eight hours later, the cells were treated with 200 nM carfilzomib (CFZ) or DMSO control for 8 hours. The cells were then harvested and subjected to quantitative RT-PCR to measure transcript levels of RUVBL1.

**Figure S2.**
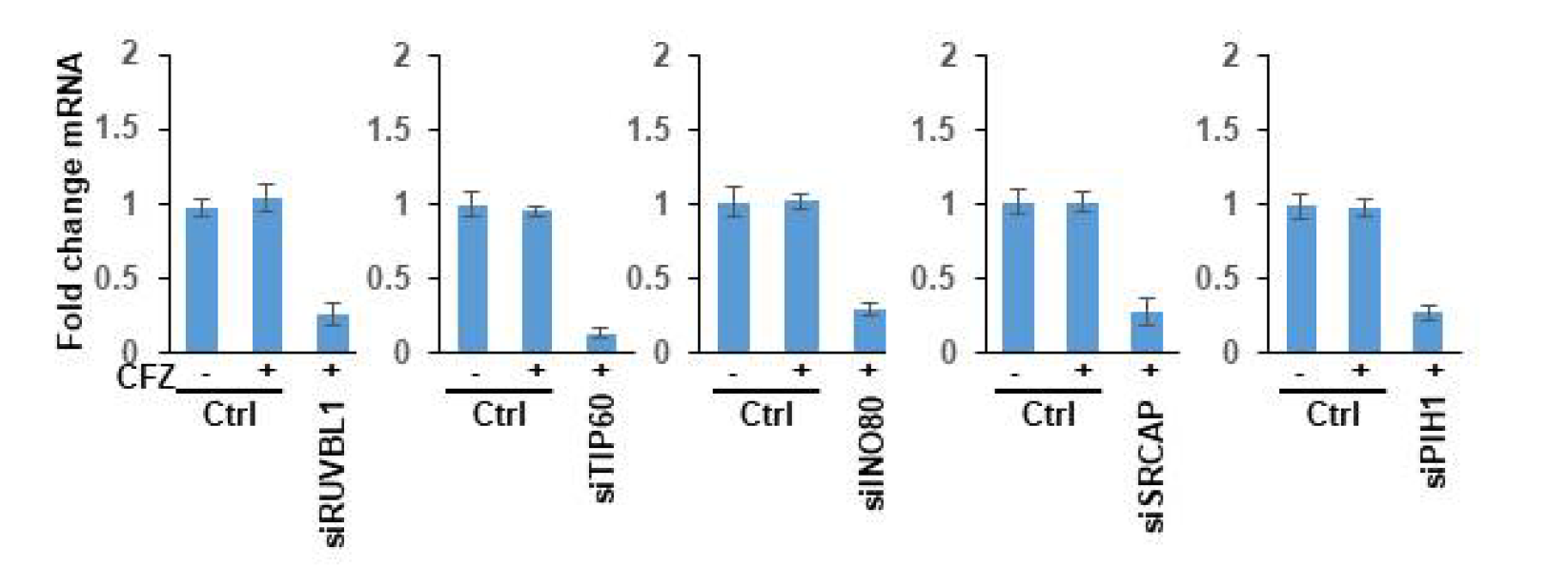
Knock-down efficiency of siRNAs targeting components of various RUVBL1-containing complexes. NIH-3T3 cells were either control transfected or with siRNAs targeting RUVBL1, TIP60, INO80, SRCAP, or PIH1. Forty-eight hours later, the cells were treated with 200 nM carfilzomib (CFZ) or DMSO control for 8 hours. The cells were then harvested and subjected to quantitative RT-PCR to measure indicated transcript levels.

**Table S1.** Results from an siRNA screen in WT 8xARE-Luc screening system.

| siRNA target | Treatment | Mean normalized Luciferase Activity | SD (n=3) |
| --- | --- | --- | --- |
| - | DMSO | 1.00 | 0.12 |
| - | Carfilzomib (CFZ) | 17.72 | 0.84 |
| DDI2 | CFZ | 3.27 | 0.33 |
| Nrf1 | CFZ | 3.17 | 0.35 |
| p97 | CFZ | 3.39 | 0.28 |
| RUVBL1 | CFZ | 3.89 | 0.35 |
| Hsp90aa1 | CFZ | 12.86 | 0.65 |
| Baz1a | CFZ | 13.19 | 0.74 |
| Baz1b | CFZ | 13.80 | 0.82 |
| Ash1l | CFZ | 13.91 | 0.56 |
| Zmynd8 | CFZ | 13.99 | 0.42 |
| Klf4 | CFZ | 14.05 | 0.36 |
| Brd1 | CFZ | 14.42 | 0.59 |
| Usp46 | CFZ | 14.51 | 0.87 |
| Brd4 | CFZ | 14.67 | 0.66 |
| Sqstm1 | CFZ | 14.82 | 0.95 |
| Hspb1 | CFZ | 14.32 | 0.39 |
| Hcfc1 | CFZ | 14.60 | 0.90 |
| Ipo11 | CFZ | 14.63 | 0.65 |
| Rbx1 | CFZ | 14.70 | 1.53 |
| Sp110 | CFZ | 14.77 | 0.88 |
| Smarca2 | CFZ | 14.87 | 1.12 |
| Sp140 | CFZ | 14.88 | 0.69 |
| Usp44 | CFZ | 14.89 | 0.37 |
| Hspa4 | CFZ | 14.99 | 0.45 |
| Trim28 | CFZ | 15.13 | 0.62 |
| Tsc1 | CFZ | 15.36 | 0.57 |
| Ubxn11 | CFZ | 15.46 | 0.64 |
| Crebbp | CFZ | 15.73 | 0.53 |
| Plk1 | CFZ | 15.80 | 1.36 |
| Brd9 | CFZ | 15.86 | 2.14 |
| Baz2a | CFZ | 15.90 | 0.65 |
| Ssrp1 | CFZ | 15.97 | 1.46 |
| Brwd3 | CFZ | 16.23 | 1.52 |
| Atad2 | CFZ | 16.32 | 0.95 |
| Brpf3 | CFZ | 16.42 | 0.35 |
| Brd7 | CFZ | 16.58 | 1.33 |
| E2f7 | CFZ | 16.66 | 1.87 |
| Hspa1a | CFZ | 16.70 | 0.54 |
| Elk1 | CFZ | 16.84 | 0.66 |
| Bptf | CFZ | 17.39 | 0.39 |
| Ep300 | CFZ | 17.75 | 0.28 |
| Fli1 | CFZ | 17.77 | 0.59 |
| Hspa2 | CFZ | 17.78 | 1.33 |
| Uba3 | CFZ | 17.79 | 2.65 |
| Cecr2 | CFZ | 17.85 | 1.95 |
| Ctcf | CFZ | 18.28 | 1.37 |
| Ewsr1 | CFZ | 18.40 | 0.87 |
| Pign | CFZ | 18.55 | 2.57 |
| Sp100 | CFZ | 18.68 | 3.56 |
| Phip | CFZ | 18.71 | 0.64 |
| Hectd1 | CFZ | 18.82 | 1.22 |
| Baz2b | CFZ | 18.92 | 0.94 |
| Brd8 | CFZ | 19.44 | 0.67 |
| Fbxo5 | CFZ | 19.49 | 1.62 |
| Dnaja2 | CFZ | 19.75 | 1.25 |
| Kmt2a | CFZ | 19.81 | 0.68 |
| Zmynd11 | CFZ | 19.91 | 1.75 |
| Kat2b | CFZ | 20.35 | 0.69 |
| Hspa8 | CFZ | 20.51 | 1.52 |
| Usp9x | CFZ | 20.69 | 0.53 |
| Trim66 | CFZ | 21.15 | 2.51 |
| Trim33 | CFZ | 21.22 | 0.39 |
| Trim24 | CFZ | 21.40 | 0.95 |
| Brd3 | CFZ | 21.51 | 3.43 |
| Capn9 | CFZ | 21.55 | 1.36 |
| Hsf1 | CFZ | 21.92 | 1.41 |
| Ube2m | CFZ | 22.32 | 2.16 |
| Smarca4 | CFZ | 22.35 | 2.40 |
| Dnajc17 | CFZ | 22.60 | 0.98 |
| Kat2a | CFZ | 22.80 | 1.43 |
| Atad2b | CFZ | 22.81 | 1.67 |
| Brwd1 | CFZ | 22.98 | 2.29 |
| Rpn1 | CFZ | 23.27 | 1.71 |
| Brdt | CFZ | 23.31 | 3.64 |
| Brd2 | CFZ | 23.53 | 1.58 |
| Bag6 | CFZ | 23.82 | 0.80 |
| Pbrm1 | CFZ | 23.92 | 1.69 |
| Brpf1 | CFZ | 24.59 | 2.87 |
| Taf1 | CFZ | 24.79 | 1.65 |
| Hsp90ab1 | CFZ | 25.09 | 2.36 |

**Table S2.**
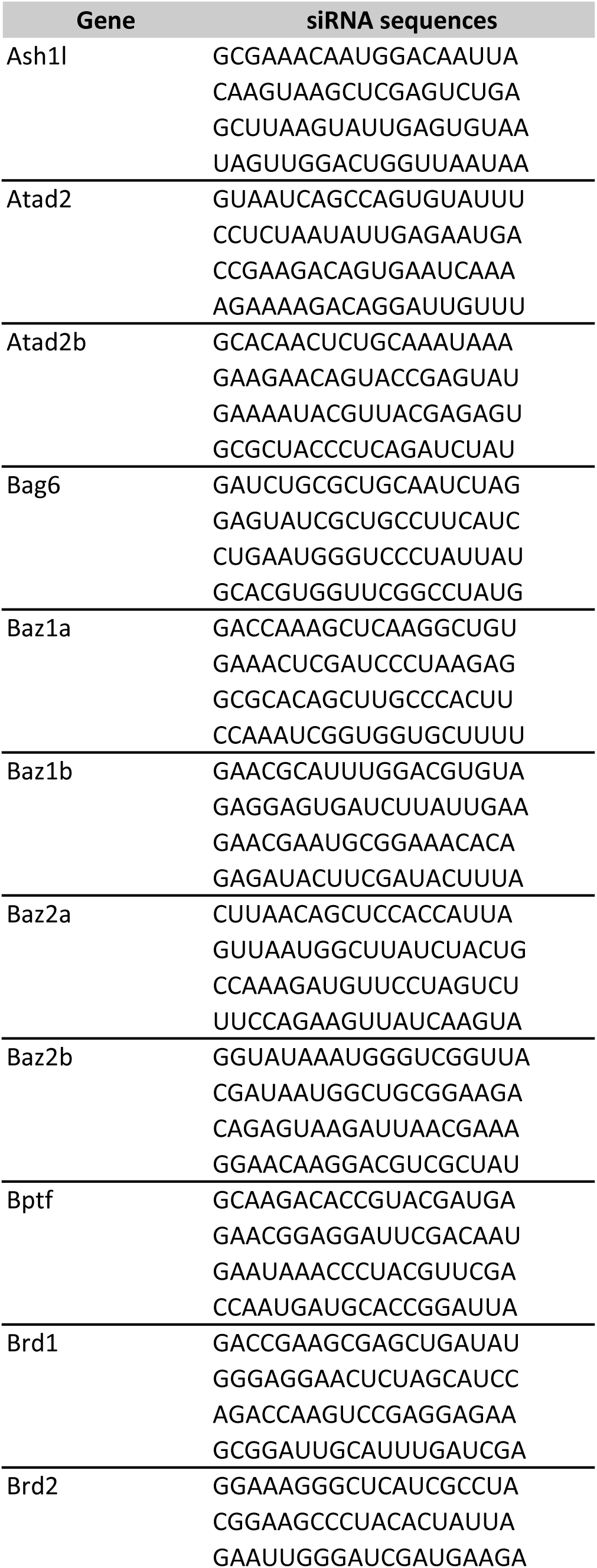
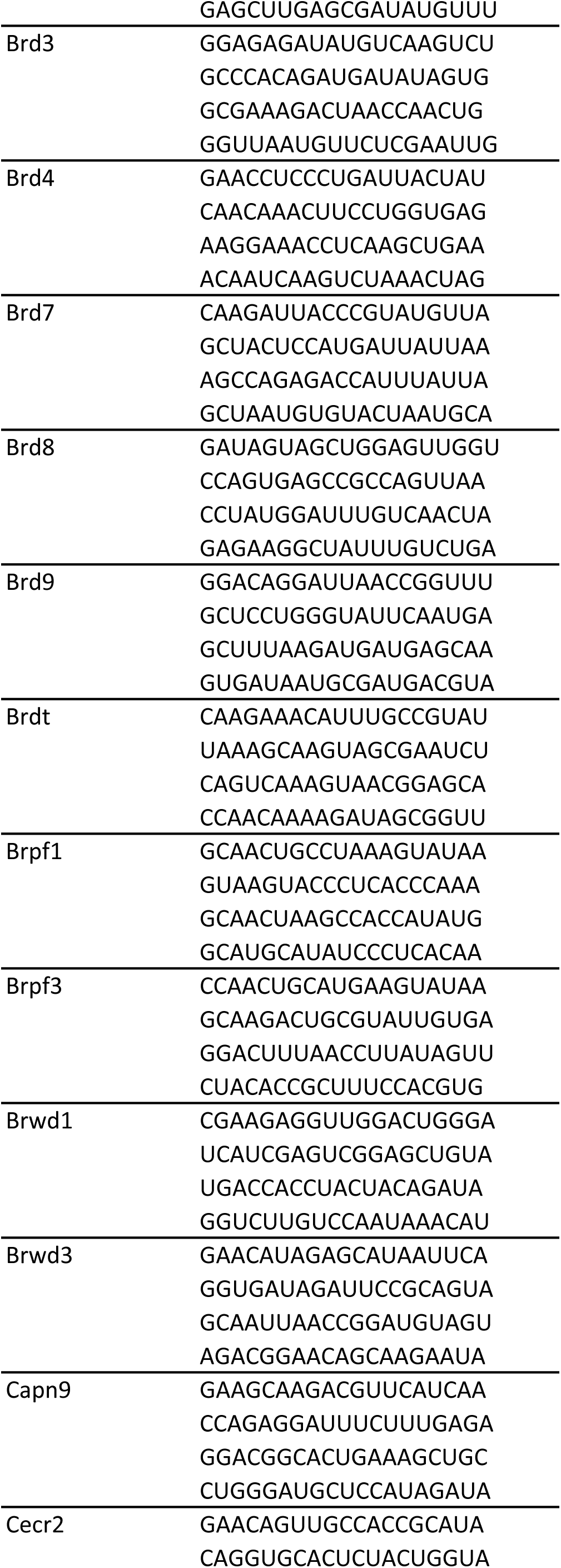
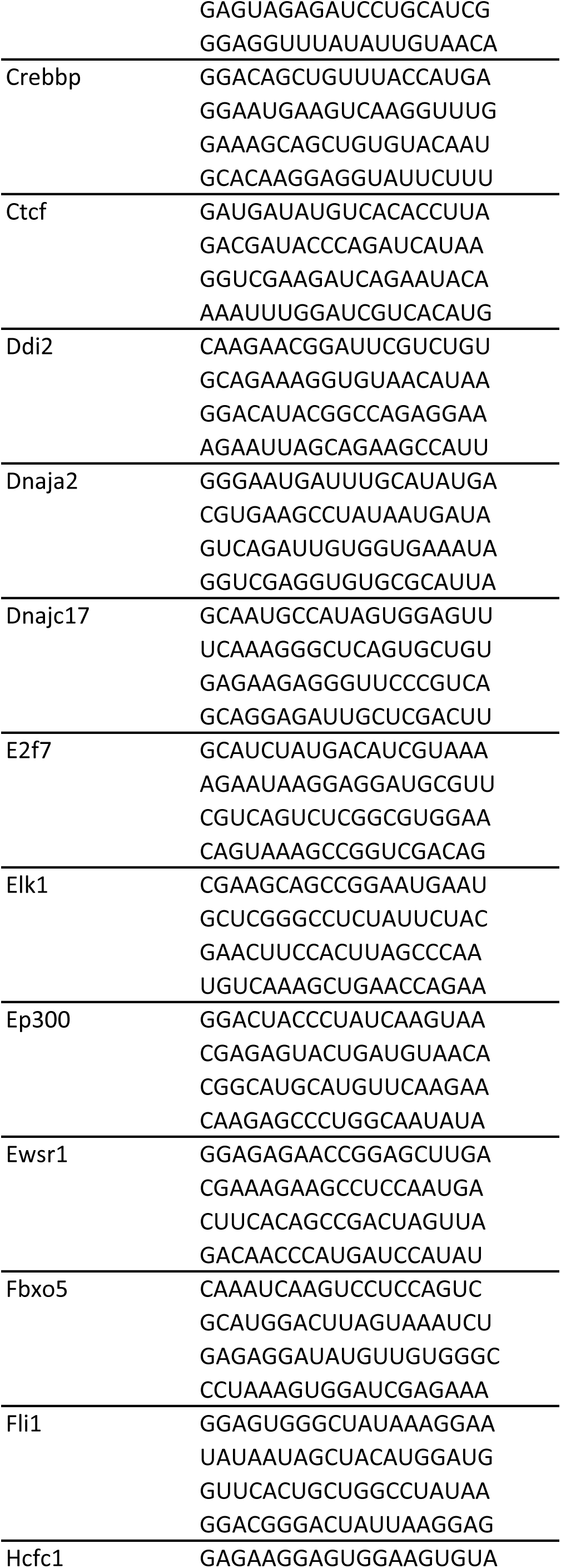
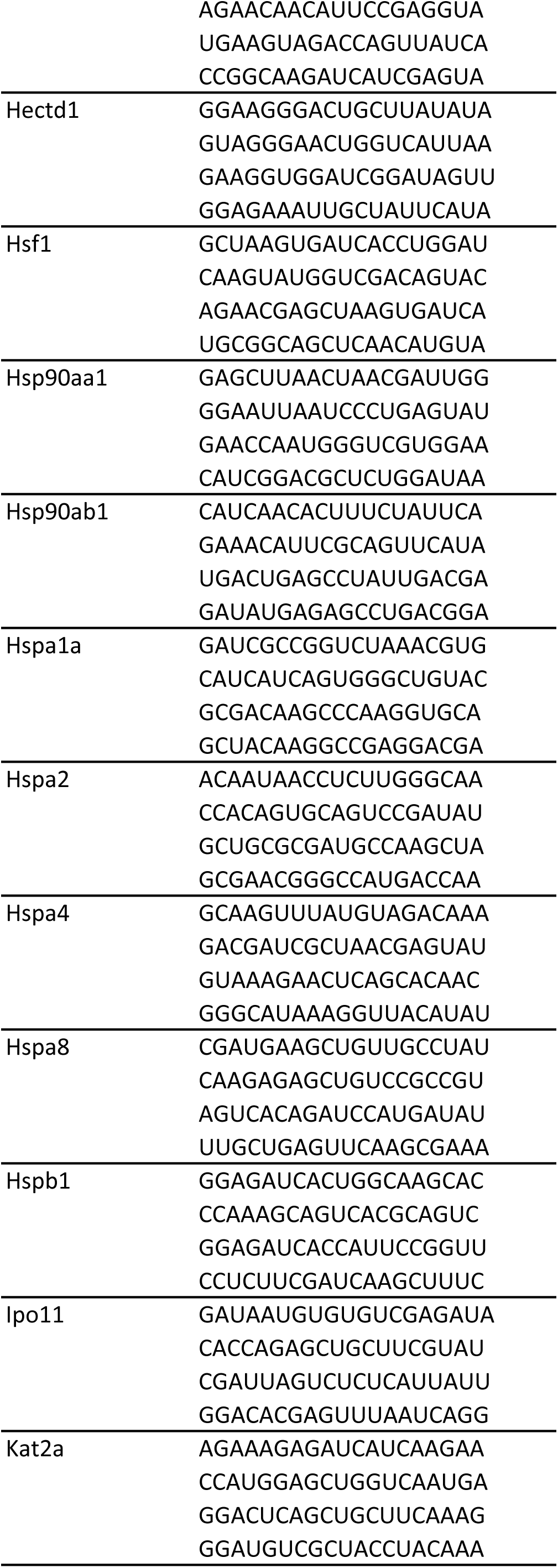
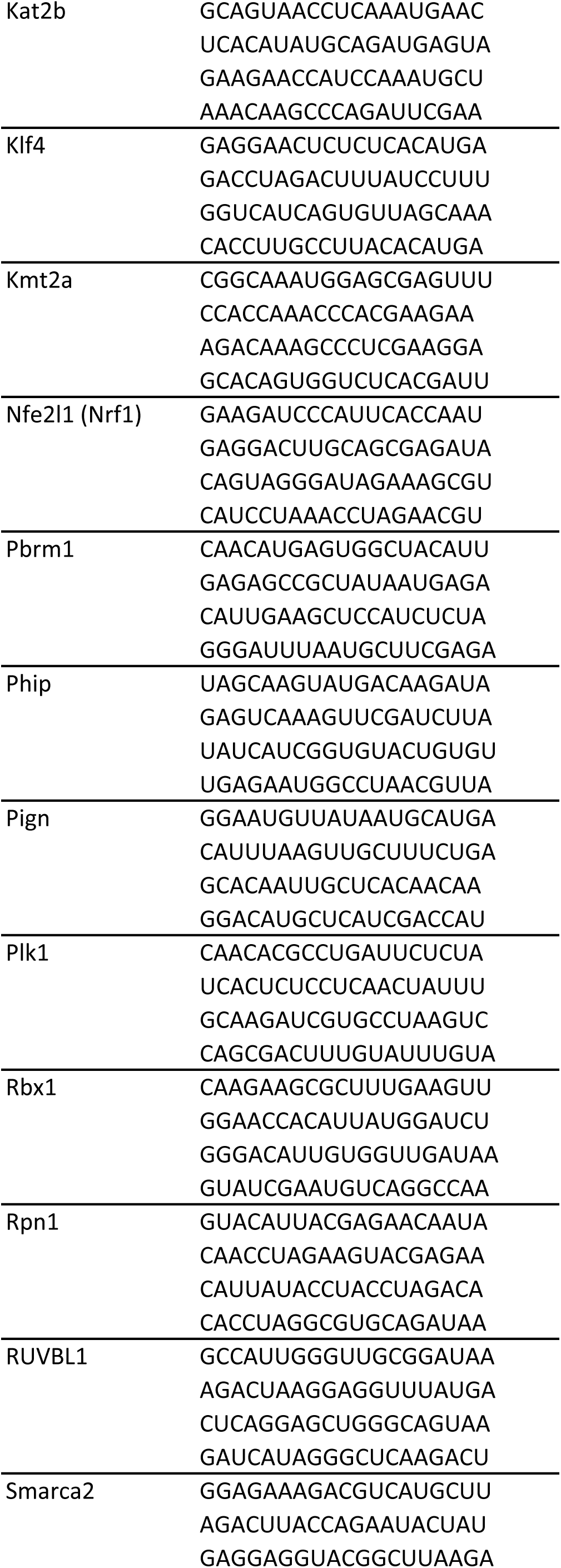
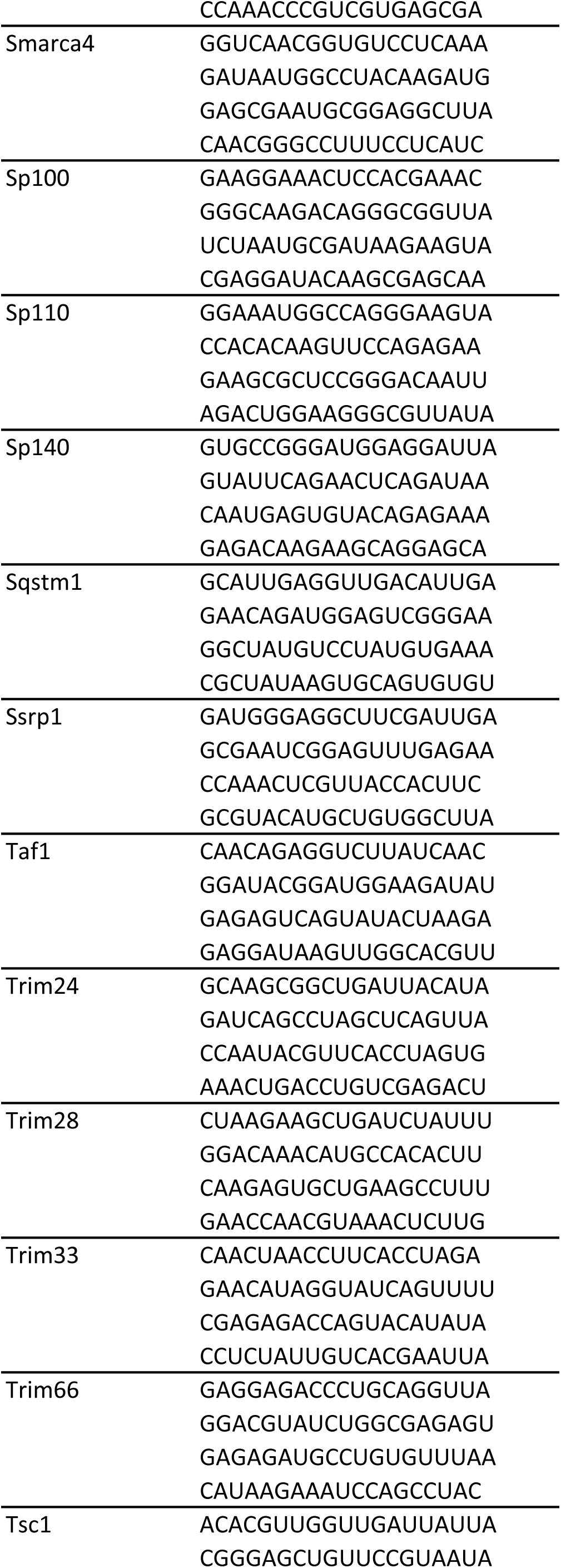
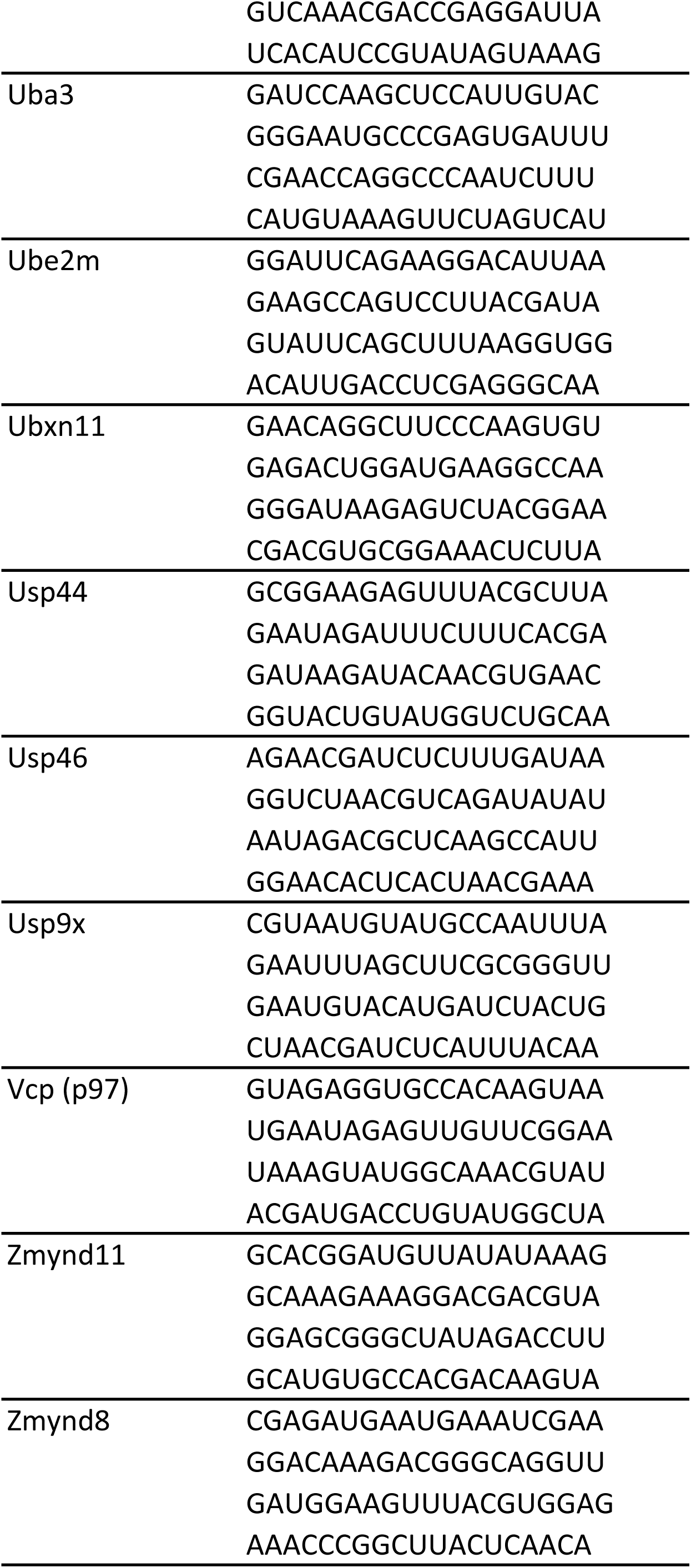
siRNA sequences for the genes targeted in the RNAi screen.

**Table S3.**
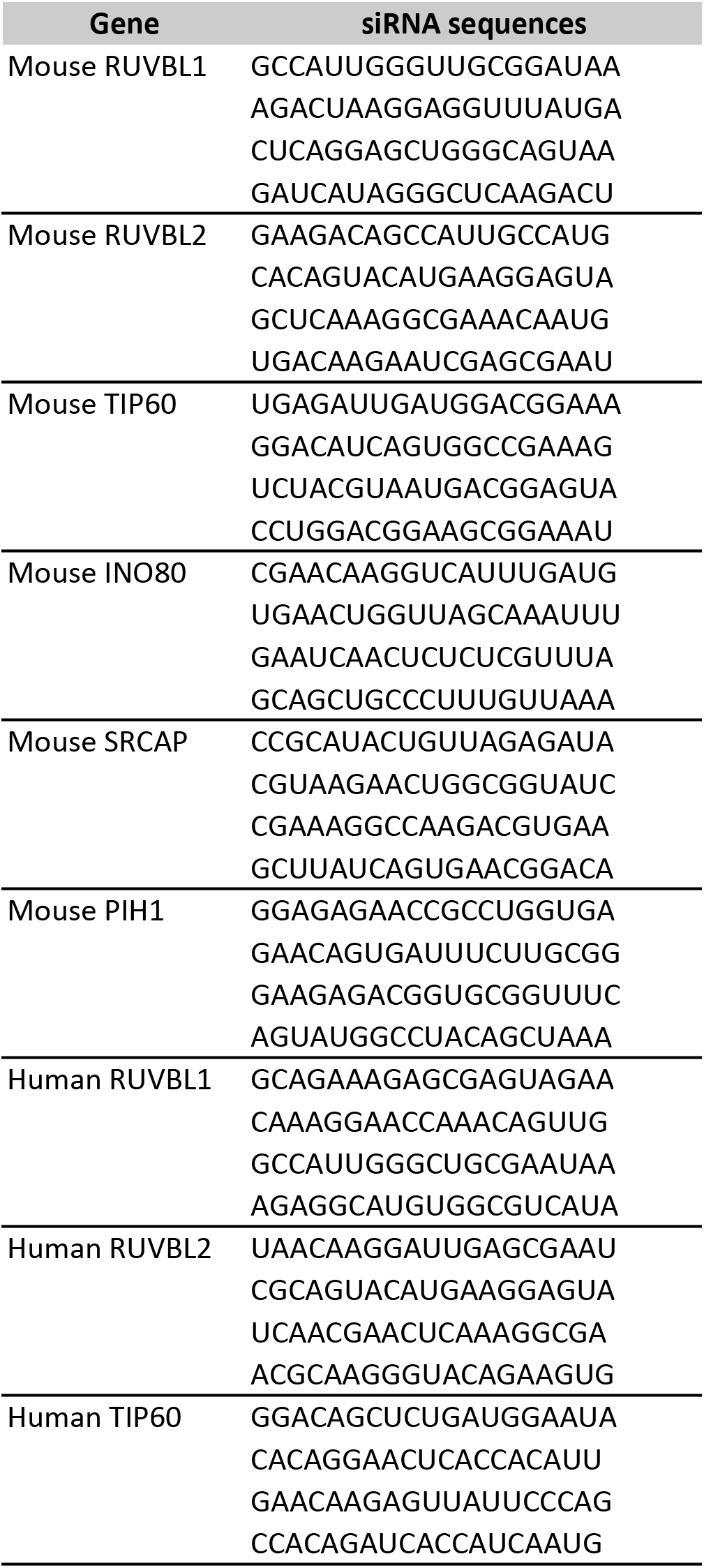
siRNA sequences for the genes targeted in experiments other than the RNAi screen.

**Table S4.**
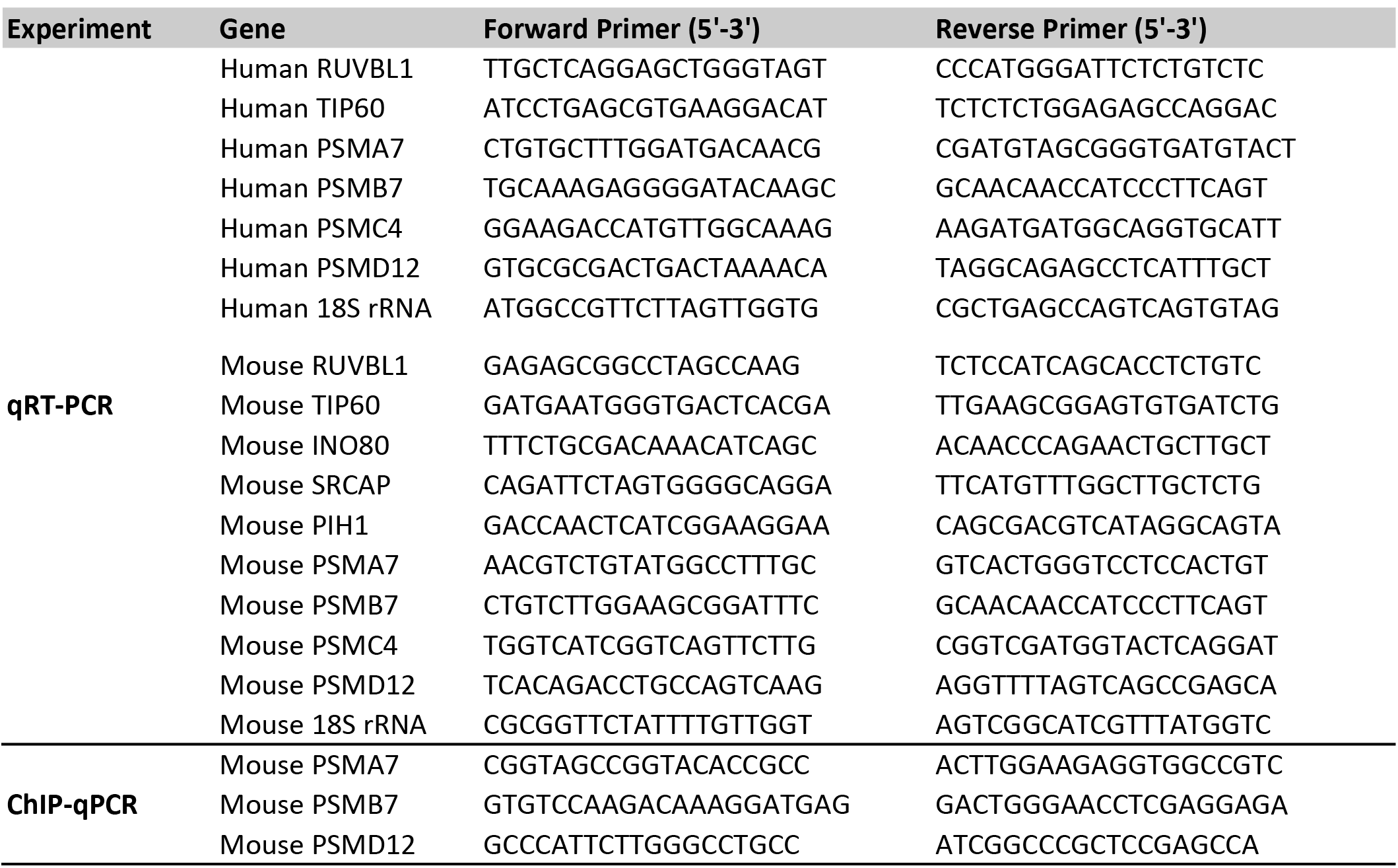
Primers used in quantitative RT-PCR (qRT-PCR) and chromatin immunoprecipitation (ChIP)-qPCR assays.

## REFERENCES

1. Lindquist, S. L., and Kelly, J. W. (2011) Chemical and biological approaches for adapting proteostasis to ameliorate protein misfolding and aggregation diseases: progress and prognosis. Cold Spring Harb Perspect Biol 3

2. Rousseau, A., and Bertolotti, A. (2018) Regulation of proteasome assembly and activity in health and disease. Nat Rev Mol Cell Biol

3. Schmidt, M., and Finley, D. (2014) Regulation of proteasome activity in health and disease. Biochim Biophys Acta 1843, 13–25

4. Radhakrishnan, S. K., Lee, C. S., Young, P., Beskow, A., Chan, J. Y., and Deshaies, R. J. (2010) Transcription factor Nrf1 mediates the proteasome recovery pathway after proteasome inhibition in mammalian cells. Mol Cell 38, 17–28

5. Sekine, H., Okazaki, K., Kato, K., Alam, M. M., Shima, H., Katsuoka, F., Tsujita, T., Suzuki, N., Kobayashi, A., Igarashi, K., Yamamoto, M., and Motohashi, H. (2018) O-GlcNAcylation Signal Mediates Proteasome Inhibitor Resistance in Cancer Cells by Stabilizing NRF1. Mol Cell Biol

6. Steffen, J., Seeger, M., Koch, A., and Kruger, E. (2010) Proteasomal degradation is transcriptionally controlled by TCF11 via an ERAD-dependent feedback loop. Mol Cell 40, 147–158

7. Tsuchiya, Y., Morita, T., Kim, M., Iemura, S., Natsume, T., Yamamoto, M., and Kobayashi, A. (2011) Dual regulation of the transcriptional activity of Nrf1 by beta-TrCP-and Hrd1-dependent degradation mechanisms. Mol Cell Biol 31, 4500–4512

8. Biswas, M., and Chan, J. Y. (2009) Role of Nrf1 in antioxidant response element-mediated gene expression and beyond. Toxicol Appl Pharmacol

9. Kim, H. M., Han, J. W., and Chan, J. Y. (2016) Nuclear Factor Erythroid-2 Like 1 (NFE2L1): Structure, function and regulation. Gene

10. Rojo de la Vega, M., Chapman, E., and Zhang, D. D. (2018) NRF2 and the Hallmarks of Cancer. Cancer Cell 34, 21–43

11. Lee, C. S., Lee, C., Hu, T., Nguyen, J. M., Zhang, J., Martin, M. V., Vawter, M. P., Huang, E. J., and Chan, J. Y. (2011) Loss of nuclear factor E2-related factor 1 in the brain leads to dysregulation of proteasome gene expression and neurodegeneration. Proc Natl Acad Sci U S A 108, 8408–8413

12. Lee, C. S., Ho, D. V., and Chan, J. Y. (2013) Nuclear factor-erythroid 2-related factor 1 regulates expression of proteasome genes in hepatocytes and protects against endoplasmic reticulum stress and steatosis in mice. FEBS J 280, 3609–3620

13. Radhakrishnan, S. K., den Besten, W., and Deshaies, R. J. (2014) p97-dependent retrotranslocation and proteolytic processing govern formation of active Nrf1 upon proteasome inhibition. Elife 3, e01856

14. Tomlin, F. M., Gerling-Driessen, U. I. M., Liu, Y.-C., Flynn, R. A., Vangala, J. R., Lentz, C. S., Clauder-Muenster, S., Jakob, P., Mueller, W. F., Ordoñez-Rueda, D., Paulsen, M., Matsui, N., Foley, D., Rafalko, A., Suzuki, T., Bogyo, M., Steinmetz, L. M., Radhakrishnan, S. K., and Bertozzi, C. R. (2017) Inhibition of NGLY1 Inactivates the Transcription Factor Nrf1 and Potentiates Proteasome Inhibitor Cytotoxicity. ACS Central Science 3, 1143–1155

15. Koizumi, S., Irie, T., Hirayama, S., Sakurai, Y., Yashiroda, H., Naguro, I., Ichijo, H., Hamazaki, J., and Murata, S. (2016) The aspartyl protease DDI2 activates Nrf1 to compensate for proteasome dysfunction. Elife 5

16. Lehrbach, N. J., and Ruvkun, G. (2016) Proteasome dysfunction triggers activation of SKN-1A/Nrf1 by the aspartic protease DDI-1. Elife 5

17. Orlowski, R. Z., and Kuhn, D. J. (2008) Proteasome inhibitors in cancer therapy: lessons from the first decade. Clin Cancer Res 14, 1649–1657

18. Le Moigne, R., Aftab, B. T., Djakovic, S., Dhimolea, E., Valle, E., Murnane, M., King, E. M., Soriano, F., Menon, M. K., Wu, Z. Y., Wong, S. T., Lee, G. J., Yao, B., Wiita, A. P., Lam, C., Rice, J., Wang, J., Chesi, M., Bergsagel, P. L., Kraus, M., Driessen, C., Kiss von Soly, S., Yakes, F. M., Wustrow, D., Shawver, L., Zhou, H. J., Martin, T. G., 3rd, Wolf, J. L., Mitsiades, C. S., Anderson, D. J., and Rolfe, M. (2017) The p97 Inhibitor CB-5083 Is a Unique Disrupter of Protein Homeostasis in Models of Multiple Myeloma. Mol Cancer Ther 16, 2375–2386

19. Vangala, J. R., Sotzny, F., Krüger, E., Deshaies, R. J., and Radhakrishnan, S. K. (2016) Nrf1 can be processed and activated in a proteasome-independent manner. Current Biology 26, R834-R835

20. Magnaghi, P., D’Alessio, R., Valsasina, B., Avanzi, N., Rizzi, S., Asa, D., Gasparri, F., Cozzi, L., Cucchi, U., Orrenius, C., Polucci, P., Ballinari, D., Perrera, C., Leone, A., Cervi, G., Casale, E., Xiao, Y., Wong, C., Anderson, D. J., Galvani, A., Donati, D., O’Brien, T., Jackson, P. K., and Isacchi, A. (2013) Covalent and allosteric inhibitors of the ATPase VCP/p97 induce cancer cell death. Nat Chem Biol 9, 548–556

21. Chatr-Aryamontri, A., Oughtred, R., Boucher, L., Rust, J., Chang, C., Kolas, N. K., O’Donnell, L., Oster, S., Theesfeld, C., Sellam, A., Stark, C., Breitkreutz, B. J., Dolinski, K., and Tyers, M. (2017) The BioGRID interaction database: 2017 update. Nucleic Acids Res 45, D369-D379

22. Ru, B., Sun, J., Tong, Y., Wong, C. N., Chandra, A., Tang, A. T. S., Chow, L. K. Y., Wun, W. L., Levitskaya, Z., and Zhang, J. (2018) CR2Cancer: a database for chromatin regulators in human cancer. Nucleic Acids Res 46, D918-D924

23. Mao, Y. Q., and Houry, W. A. (2017) The Role of Pontin and Reptin in Cellular Physiology and Cancer Etiology. Front Mol Biosci 4, 58

24. Venteicher, A. S., Meng, Z., Mason, P. J., Veenstra, T. D., and Artandi, S. E. (2008) Identification of ATPases pontin and reptin as telomerase components essential for holoenzyme assembly. Cell 132, 945–957

25. Izumi, N., Yamashita, A., Iwamatsu, A., Kurata, R., Nakamura, H., Saari, B., Hirano, H., Anderson, P., and Ohno, S. (2010) AAA+ proteins RUVBL1 and RUVBL2 coordinate PIKK activity and function in nonsense-mediated mRNA decay. Sci Signal 3, ra27

26. Rajendra, E., Garaycoechea, J. I., Patel, K. J., and Passmore, L. A. (2014) Abundance of the Fanconi anaemia core complex is regulated by the RuvBL1 and RuvBL2 AAA+ ATPases. Nucleic Acids Res 42, 13736–13748

27. Gnatovskiy, L., Mita, P., and Levy, D. E. (2013) The human RVB complex is required for efficient transcription of type I interferon-stimulated genes. Mol Cell Biol 33, 3817–3825

28. Jha, S., Gupta, A., Dar, A., and Dutta, A. (2013) RVBs are required for assembling a functional TIP60 complex. Mol Cell Biol 33, 1164–1174

29. Frankland-Searby, S., and Bhaumik, S. R. (2012) The 26S proteasome complex: an attractive target for cancer therapy. Biochim Biophys Acta 1825, 64–76

30. Petrocca, F., Altschuler, G., Tan, S. M., Mendillo, M. L., Yan, H., Jerry, D. J., Kung, A. L., Hide, W., Ince, T. A., and Lieberman, J. (2013) A genome-wide siRNA screen identifies proteasome addiction as a vulnerability of basal-like triple-negative breast cancer cells. Cancer Cell 24, 182–196

31. Tarangelo, A., Lo, N., Teng, R., Kim, E., Le, L., Watson, D., Furth, E. E., Raman, P., Ehmer, U., and Viatour, P. (2015) Recruitment of Pontin/Reptin by E2f1 amplifies E2f transcriptional response during cancer progression. Nat Commun 6, 10028

32. Sapountzi, V., Logan, I. R., and Robson, C. N. (2006) Cellular functions of TIP60. Int J Biochem Cell Biol 38, 1496–1509

33. Perez-Perri, J. I., Dengler, V. L., Audetat, K. A., Pandey, A., Bonner, E. A., Urh, M., Mendez, J., Daniels, D. L., Wappner, P., Galbraith, M. D., and Espinosa, J. M. (2016) The TIP60 Complex Is a Conserved Coactivator of HIF1A. Cell Rep 16, 37–47

34. Gaughan, L., Logan, I. R., Cook, S., Neal, D. E., and Robson, C. N. (2002) Tip60 and histone deacetylase 1 regulate androgen receptor activity through changes to the acetylation status of the receptor. J Biol Chem 277, 25904–25913

35. Patel, J. H., Du, Y., Ard, P. G., Phillips, C., Carella, B., Chen, C. J., Rakowski, C., Chatterjee, C., Lieberman, P. M., Lane, W. S., Blobel, G. A., and McMahon, S. B. (2004) The c-MYC oncoprotein is a substrate of the acetyltransferases hGCN5/PCAF and TIP60. Mol Cell Biol 24, 10826–10834

36. Elkaim, J., Castroviejo, M., Bennani, D., Taouji, S., Allain, N., Laguerre, M., Rosenbaum, J., Dessolin, J., and Lestienne, P. (2012) First identification of small-molecule inhibitors of Pontin by combining virtual screening and enzymatic assay. Biochem J 443, 549–559

37. Elkaim, J., Lamblin, M., Laguerre, M., Rosenbaum, J., Lestienne, P., Eloy, L., Cresteil, T., Felpin, F. X., and Dessolin, J. (2014) Design, synthesis and biological evaluation of Pontin ATPase inhibitors through a molecular docking approach. Bioorg Med Chem Lett 24, 2512–2516

38. Gao, C., Bourke, E., Scobie, M., Famme, M. A., Koolmeister, T., Helleday, T., Eriksson, L. A., Lowndes, N. F., and Brown, J. A. (2014) Rational design and validation of a Tip60 histone acetyltransferase inhibitor. Sci Rep 4, 5372

39. Roeten, M. S. F., Cloos, J., and Jansen, G. (2018) Positioning of proteasome inhibitors in therapy of solid malignancies. Cancer Chemother Pharmacol 81, 227–243

40. Levy, C., Khaled, M., Iliopoulos, D., Janas, M. M., Schubert, S., Pinner, S., Chen, P. H., Li, S., Fletcher, A. L., Yokoyama, S., Scott, K. L., Garraway, L. A., Song, J. S., Granter, S. R., Turley, S. J., Fisher, D. E., and Novina, C. D. (2010) Intronic miR-211 assumes the tumor suppressive function of its host gene in melanoma. Mol Cell 40, 841–849

